# Effects of Ca^2+^ on the Structure and Dynamics of PIP3 in Model Membranes Containing PC and PS

**DOI:** 10.1101/2024.05.28.596302

**Authors:** Ashley D. Bernstein, Yanxing Yang, Thomas M. Osborn Popp, Gertrude Asante Ampadu, Gobin Raj Acharya, Andrew J. Nieuwkoop

**Affiliations:** Department of Chemistry and Chemical Biology, Rutgers, The State University of New Jersey, Piscataway, New Jersey 08854, United States

**Keywords:** Lipids, Solid-state NMR, NMR, PIP3, PS, Ca^2+^

## Abstract

Phosphatidylinositol phosphates (PIPs) are a family of seven different eukaryotic membrane lipids that have a large role in cell viability, despite their minor concentration in eukaryotic cellular membranes. PIPs tightly regulate cellular processes such as cellular growth, metabolism, immunity, and development through direct interactions with partner proteins. Understanding the biophysical properties of PIPs in the complex membrane environment is important to understand how PIPs selectively regulate a partner protein. Here we investigate the structure and dynamics of PIP3 in lipid bilayers that are simplified models of the natural membrane environment. We probe the effects of the anionic lipid phosphatidylserine (PS) and the divalent cation Ca^2+^. We use solution and solid-state ^1^H, ^31^P, and ^13^C NMR all at natural abundance combined with MD simulations to characterize the structure and dynamics of PIPs. ^1^H and ^31^P 1D spectra show good resolution at high temperatures with isolated peaks in the headgroup, interfacial, and bilayer regions. Site specific assignment of these 1D reporters were made and used to measure the effects of Ca^2+^ and PS. In particular, the resolved ^31^P signals of the PIP3 headgroup allowed for extremely well localized information about PIP3 phosphate dynamics, which the MD simulations were able to help explain. Cross polarization kinetics provided additional site-specific dynamics measurements for the PIP3 headgroups.

## INTRODUCTION

Phosphatidylinositol phosphates (PIPs) are a family of seven different eukaryotic membrane lipids that result from the phosphorylation of the lipid phosphatidylinositol (PI)’s head group once (PI(3)P, PI(4)P, and PI(5)P), twice (PI(3,4)P2, PI(3,5)P2, and PI(4,5)P2), or three times (PIP3; PI(3,4,5)P3).^1–4^ Despite their minor concentration (less than 5%) in eukaryotic cellular membranes, PIPs play a significant role in regulating cellular processes such as cellular growth, metabolism, immunity, and development through interactions with proteins.^1–3^ Activation of PIP binding proteins is often specific to a single PIP species, despite the apparent similarity between PIPs of differing extents of phosphorylation.^1–4^. By directly measuring the structure and dynamics of PIPs in phospholipid membranes, we aim to discover potential molecular mechanisms by which PIPs may be discriminated, with implications for understanding how proteins achieve specific binding to a particular PIP species.

In this work, we focus on understanding PIP3 in model phospholipid membranes to elucidate the structural and dynamic properties of this lipid in the context of the cellular membrane environment. PIP3 is found primarily on the cytosolic side of the cellular membrane and activates proteins that control a variety of processes, including cellular growth, cell apoptosis, and insulin signaling. Mis-regulation of PIP3 pathways can lead to obesity, diabetes, and other metabolic diseases. ^5–8^ The molecular mechanisms of PIP binding likely involve additional interactions with the bilayer, and thus our investigations of PIP3 are carried out in lipid bilayers that are simplified models of the natural cytosolic membrane environment to systematically delineate the role of each component. Here we will focus on two, the divalent cation Ca^2+^ and the anionic phospholipid phosphatidylserine (PS). Ca^2+^ exists in low μM concentrations in the intracellular environment, with local increases in concentration serving a signaling role.^9^ For example, C2 domains bind PIPs in a Ca^2+^ dependent manner, and anionic lipids, such as PIP2, have been shown to cluster in the presence of Ca^2+^.^10–15^ Our group has investigated the effects of Ca^2+^ on the binding of the F0 and PH domains of the human cell adhesion regulator kindlin-2 to PIP2 and PIP3.^16^ While up to 50% of the lipids present in eukaryotic membranes have phosphatidylcholine (PC) as the headgroup,^5–7^ PS is present in abundance in the cytosolic membrane leaflet^5–7, 17^ PS has been shown to be essential for the activation of PIP-binding proteins such as the PH domain of the PIP3-binding protein Akt.^17^. We seek to understand the mode of action of these positive and negatively charged modulators of PIP3 activity.

The primary experimental technique used in this study is magic angle spinning (MAS) solid-state NMR (ssNMR). MAS ssNMR is a premier technique for studying systems that lack long range order, such as fibrils, large membrane proteins, and lipids.^18–22^ ssNMR is able to provide atomic resolution information on these relatively complicated systems that are difficult to study with other techniques. This study focuses on ^31^P and ^1^H as probes for our lipid system, as both are highly sensitive NMR nuclei naturally present in key areas of the PIP3 without the need for any isotopic or chemical labeling. ^31^P is particularly helpful to study PIPs – ^31^P is located on the head group of PIPs in addition to being present in the interfacial region for all phospholipids (Figure 1a and b). Similarly, protons are abundant in lipids throughout their structure (Figure 1a and b), with ^1^H sites with distinct chemical shifts present on the head group, interfacial glycerol group, and the fatty acid tails. The samples in this work are analyzed by 1D ^31^P and ^1^H spectra, static 1D ^31^P spectra, 2D HP and HC correlations, and T_1_ and T_2_ relaxation experiments for ^31^P. In addition, molecular dynamics (MD) simulations were employed to provide atomic insights of the conformation and dynamics of PIP3. The highly mobile membrane mimetic (HMMM) method was used to increase sampling of both the lateral and rotational dynamics of lipids within a feasible time scale. We are able to probe the structure and dynamics of PIP3, as well as PC and PS, in model lipid bilayers and receive distinct information from the headgroup, interfacial, and tail regions. Our results show significant variations in lipid conformation and dynamics upon adding Ca^2+^, with alterations in hydrogen bonding patterns and decreases in lateral diffusion and headgroup rotations.

**Figure 1.**
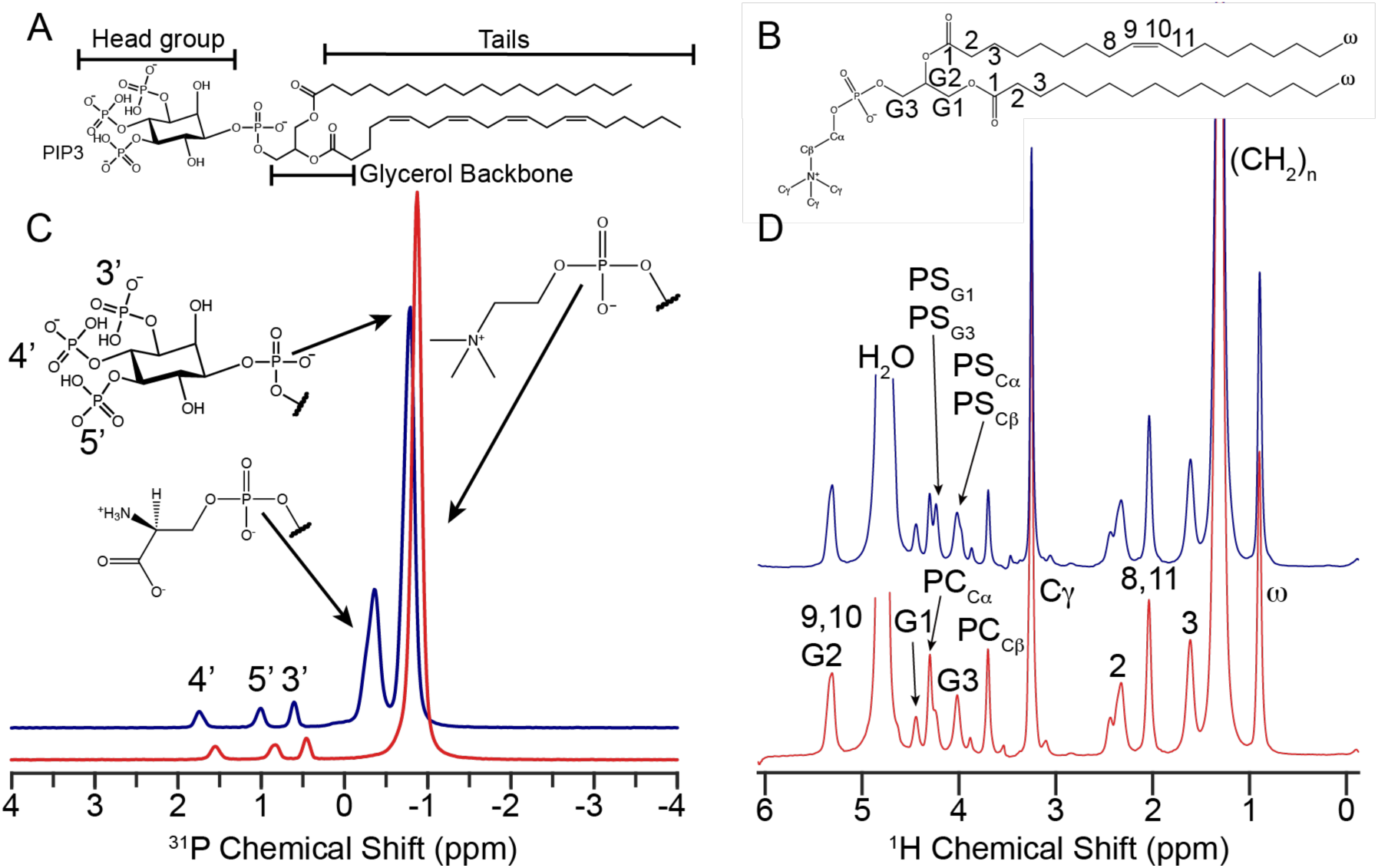
Structure and chemical shift assignment of lipids used in this study. Lipid structures of A. PIP3 and B. POPC and 1D Spectra and assignment of 5:95 PIP3/PC (red) and 5:70:25 PIP3/PC/PS (navy) liposomes C. ^31^P 1D and D. ^1^H 1D. Spectra were acquired at 20 °C and 15 kHz MAS on a 600 MHz spectrometer. Each phosphate present on the PIP3 head group is resolvable from each other, allowing for site specific analysis.

## RESULTS AND DISCUSSION

To understand the behavior of PIP3 (Figure 1A) in the inner leaflet of the plasma membrane, we choose liposomes as our model membrane. Liposomes are spherical lipid bilayers which are relatively simple to make and have easily tunable composition and size, making them excellent membrane models.^23, 24^ Liposomes were made with two lipid compositions, 5:95 PIP3/POPC (hereafter PIP3/PC) and 5:25:70 PIP3/POPS/POPC (hereafter PIP3/PC/PS). The 5% concentration of PIP3 mimics the higher end of the physiologically relevant concentration of PIP3, which helps with sensitivity of our NMR experiments. Similarly, 25% is a good representation of the inner leaflet concentration of PS, while the 70% PC stands in for all the zwitterionic components (PC, PE and SM primarily). For both PC and PS the PO tail arraignment also provides a more realistic lipid system than the less complicated DO tails sometimes used in biophysical studies, making our samples appropriate for the investigation of bulk lipid dynamics. ^31^P 1D spectrum of PIP3/PC (Figure 1C, red) show well resolved signals for the 3’, 4’, and 5’ phosphates on the PIP3 head group, while the stronger peak at -0.8 ppm represents the glycerol phosphates of PC and PIP3, which have the same chemical shift.^22^ The ^1^H 1D (Figure 1D, red) shows resolved signals for protons bound to the carbons of the headgroup (Cα, Cβ, Cγ), interfacial glycerols (G1-3), and tail regions (1-17, with the terminal methyl group labeled as omega) of POPC. The ^31^P 1D of PIP3/PC/PS can be seen in Figure 1C (blue). The additional peak at -0.36 ppm is assigned to the glycerol phosphate of PS.^25^ The PIP3 inositol phosphate peaks show narrower linewidths, especially for the 4’ site, and a shift to higher (downfield) chemical shifts, which represents deprotonation of the phosphate group, due to the presence of negatively charged PS. The ^1^H 1D (Figure 1D) shows no obvious changes to the tail region (0-3ppm) of the spectra, as might be expected given the identical structure of the tails. In the head group region (3-6 ppm) the head group protons of PS with different ^1^H chemical shifts can be identified. Assignments in this more congested region of the spectrum were aided with the use of 2D spectra (Figure 2). The slight increase in resolution indicates an increase in motion for the PIPs. Perhaps due to reduced direct interactions with the headgroup of PC which we will describe more in detail below.

**Figure 2.**
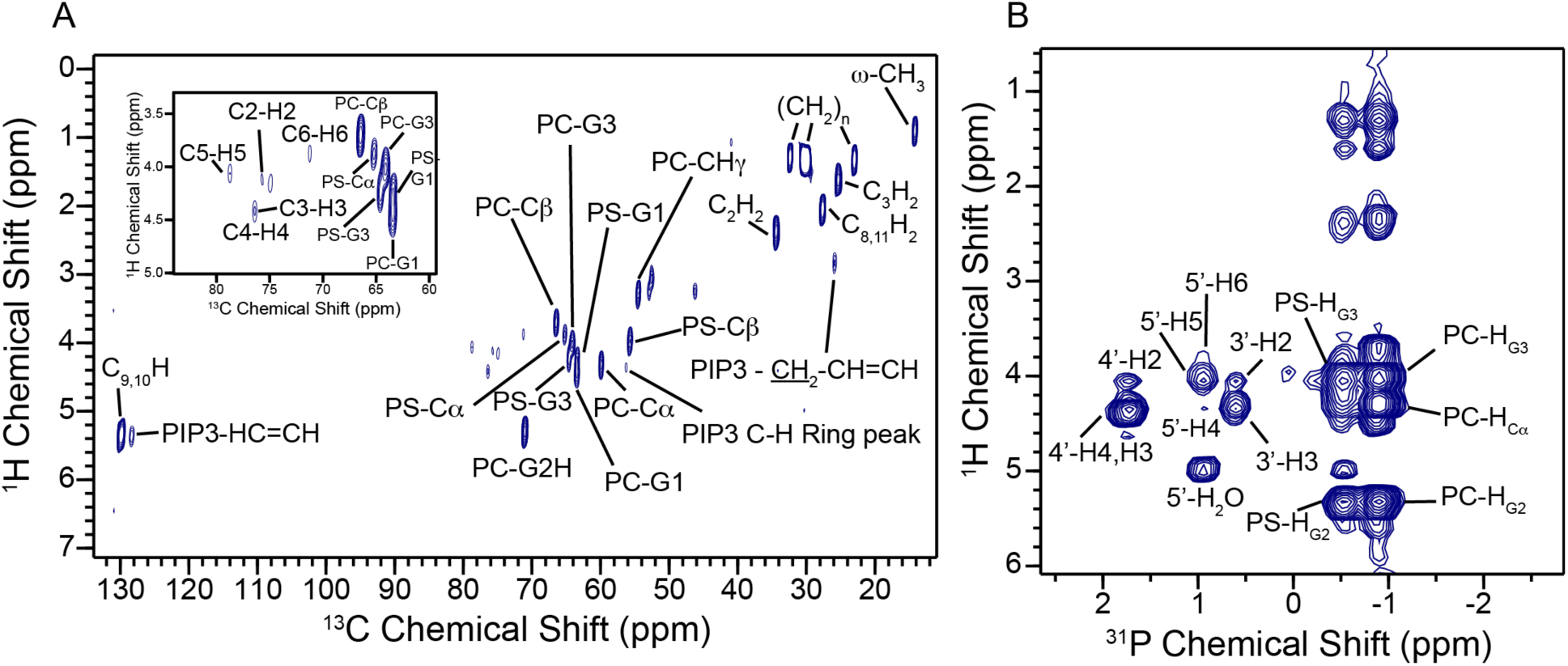
Assignment of 2D heteronuclear spectra for 5:70:25 PIP3/PC/PS liposomes. A. HC INEPT assignment of PIP3/PC/PS liposomes with inset on the head group region showing PIP3 inositol ring assignment. B. HP CP 2D spectrum of PIP3/PC/PS liposomes with assignment.

Chemical shift assignments for PIP3, PC and PS were performed using 2D heteronuclear correlation experiments on PIP3/PC/PS liposomes (Figure 2) These spectra also helped resolve PIP3 inositol peaks in the ^1^H dimension which were obscured by the water signal and stronger PC and PS peaks in 1D spectra. The HC 2D spectrum (Figure 2A) was acquired at 45 °C, well above the phase transition for PC/PS, resulting in high mobility, allowing the acquisition of a solution-like through bond INEPT experiment. Conversely, the HP 2D spectra in Figure 2B was acquired at -5 °C, below the phase transition, using through-space cross polarization experiments which require more rigid head groups to see correlations to the phosphate groups which have no directly bound protons. In addition to expected correlations based on chemical structure, some notable cross peaks appear which provide information beyond just assignments. For example a peak at the ^31^P chemical shifts of PS and the 5’ PIP3 phosphate and the ^1^H chemical shift of water which are the result of protons near the phosphate groups exchange with bulk H_2_O.^21^ Additionally, ^31^P-^1^H cross peaks appear between the glycerol phosphate of PC and PS and proton sites in the lipid tail. These correlations represent distances that are much longer than are possible for a direct transfer if the tails are extended and must be due to ^1^H-^1^H spin diffusion during the long contact times used here or disordered lipid bilayers which allow for sharply bent lipid tails. We take advantage of this reporter of the tails in our analysis. Solution NMR of PC/PS liposomes and the soluble PIP3 head group analogue Inositol 1,3,4,5-tetrakisphosphate (IP4) were also acquired. (Figure S1 and S2) As expected, some signals from the interfacial region were weak due to restriction of motion and the size of the liposomes used. Our IP4 spectra agree with the previous assignments.^26^ PIP3 has one polyunsaturated tail, unlike the mono-unsaturation in POPC and POPS, so some tail resonances of PIP3 are resolvable from PC and PS and are assigned. The only ambiguity at this time are a second set of peaks near the PC Cγ-H3 peak which occur in the presence of PS. We hypothesize these signals come from choline involved in a salt bridge with the negatively charged PS, something we observe in our MD simulations, breaking the symmetry of the 3 Cγ sites and resulting in the three additional peaks.

To measure the effects of Ca^2+^ on the inter and intramolecular dynamics of PIP3 and the bilayer as a whole, PIP3/PC samples with Ca^2+^ at concentrations (0 M, 0.1 μM, 1 μM, 5 μM, 10 μM, and 100 μM) spanning the resting intercellular calcium concentration (low μM) were made. In the presence of Ca^2+^, the linewidths of the PIP3 head group peaks are broader (Figure 3A), even at the lowest concentration of 0.1 μM. Similar to what was observed from the addition of PS, the PIP3 head group peaks are shifted slightly downfield, a phenomenon also observed in PI(4,5)P2^27^. This change in both lineshape and chemical shift are indicative of strong interactions between the PIP3 head group phosphates and Ca^2+^ and point to Ca^2+^ restricting the motion of the lipids. This view is supported by our MD simulations which show all Ca^2+^ ions in our simulations immediately sequestered by a phosphate group at the start of each simulation. Once there is one Ca^2+^ for each PIP3 pair (100 μM), there is a marked shift in the spectrum. We hypothesize this is the result of each PIP3 forming ion-mediated dimers, or higher order structures, like those observed in our MD simulations (Figure 3D). To better understand if the broadening of lines in our NMR experiments is the result of reduction of motion or an increase in sample heterogeneity, NMR relaxation experiments were performed. By measuring the T_1_ and T_2_ relaxation times for each ^31^P signal in our spectra, these experiments give site-specific information on the dynamics of each lipid species present. The mechanisms of T_1_ and T_2_ relaxation are sensitive to motions in ps-ns timescales, while T_2_ is also affected by slower exchange events as well as overall sample heterogeneity. The ^31^P T_2_ values (Figure 3B) for all three PIP3 head group phosphates drop significantly in the 0.1 μM Ca^2+^ PC/PIP3 sample, as expected from the ^31^P 1D spectra. From 1 μM to 100 μM a clear trend of decreasing T_2_ with increasing calcium concentration is observed for each PIP3 head group phosphate, with the 4’ having the lowest value, and the 3’ the highest value at each concentration perhaps as a result of the dimer structure shown in Figure 3D which allows the most freedom of motion to the 3’ phosphate. The T_2_ of the glycerol phosphates of PC and PIP3 do not exhibit a large change after the initial drop until 100 μM where an increase is seen. This is consistent with all PIP3 having dimerized, decreasing PIP3 interactions with PC and improving bilayer homogeneity. The ^31^P T_1_ values (Figure 3C) show an initial decrease for all ^31^P species at 0.1 μM Ca^2+^ but then remain constant at higher [Ca^2+^]. This is especially true for the PC ^31^P T_1_ which has the least change upon Ca^2+^ addition and changes the least throughout thereafter, while the ^31^P PIP3 head group phosphates have more variability but oscillate about the same ^31^P T_1_ values. The drop in ^31^P T_2_ combined with the lack of change for the ^31^P T_1_ indicates that higher concentrations of Ca^2+^ result in changes in dynamics at slower time scales, consistent with changes in lateral diffusion of lipids within the bilayer. This conclusion is based on assigning the motions in the ps-ns range to the free rotation around a bond necessary for phosphate group rotations, and the slower ms timescale motions we assign to long-range lateral diffusion on a nM scale liposomes.

**Figure 3.**
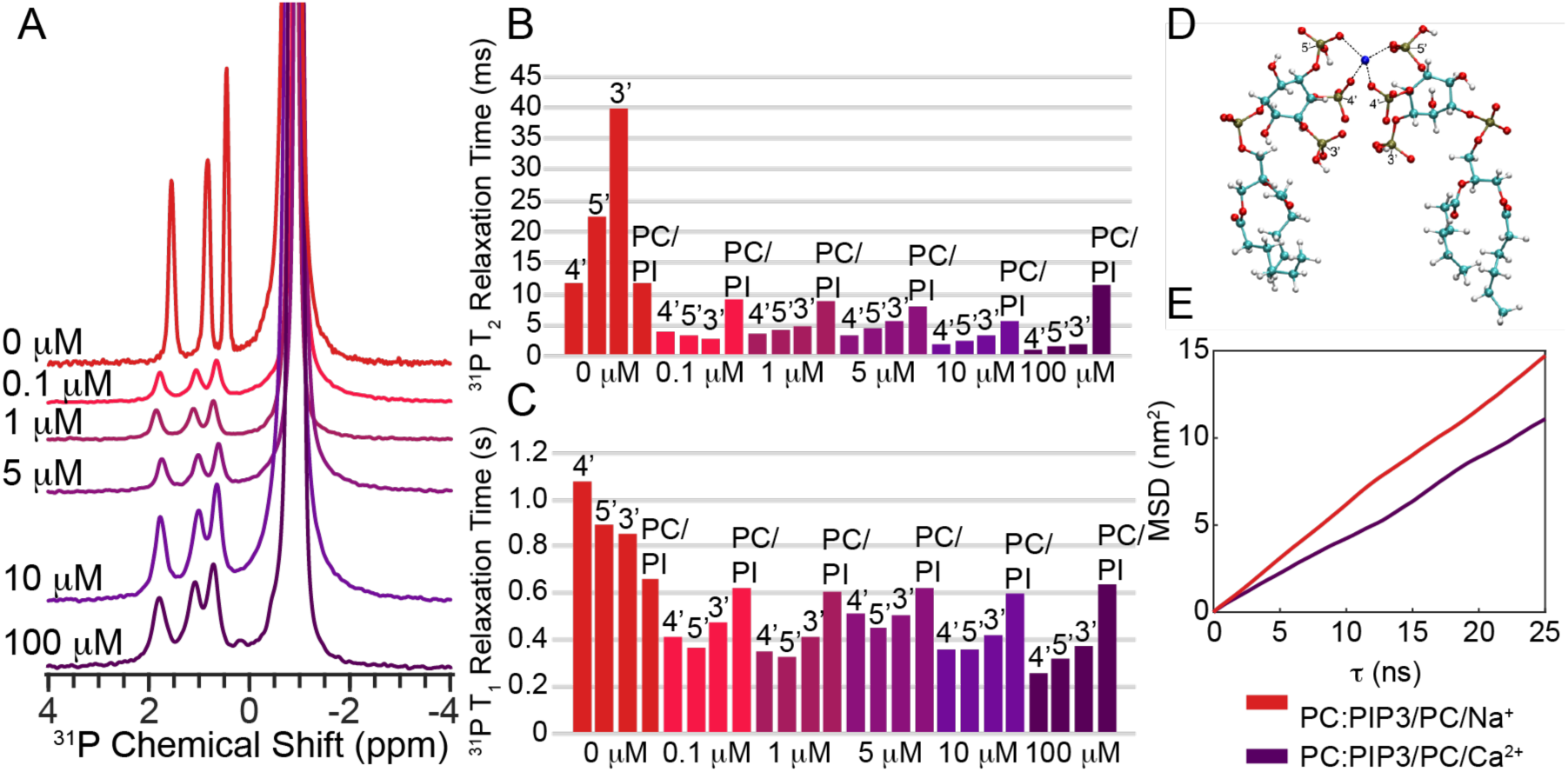
^31^P and MD simulations/lateral diffusion data for PIP3/PC liposomes at different Ca^2+^ concentrations from 0 μM-100 μM. A. ^31^P 1D spectra of PIP3/PC liposomes at each Ca^2+^ concentration. Changes in chemical shift, peak intensity, line width, and resolution are seen. B. Site specific ^31^P T_2_ relaxation data shows a large initial drop in T_2_ relaxation time for all phosphates on the PIP3 head group upon Ca^2+^ addition, which is also seen in the C. site specific ^31^P T_1_ relaxation data. D. A cartoon representation of two PIP3 headgroups interacting and forming a dimer in the presence of Ca^2+^ with distances. E. The MD lateral diffusion data for PC within the PIP3/PC liposomes with Na^+^ (red) and Ca^2+^ (purple).

To provide additional insight into the effects of calcium on lipid headgroup dynamics and lateral diffusion we performed all atom MD simulations using the Amber17 force field. Because lateral diffusion of lipids requires timescales far beyond those available using traditional MD we utilized the highly mobile membrane memetic (HMMM) strategy. This method constructs a normal bilayer and then truncates all the lipid tails after 6 carbons, allowing them to freely float on an organic solvent to increase their ability to diffuse past each other. PIP3 dimers form rapidly in these simulations (Figure 3D). Larger PIP3 trimers and tetramers were also observed. Measurements of lateral diffusion taken from the last 10 ns of 100 ns simulations of PIP3/PC bilayers (Figure 3E) show a moderate decrease in PC diffusion, supporting our interpretation of the phosphorous relaxation data.

The structures observed in MD and changes in motion implied by the T_1_ and T_2_ relaxation data led us to desire a more direct measurement of motion for the individual phosphate groups in our samples. The HP 2D experiments (Figure 2B) use cross polarization for the ^1^H to ^31^P transfer. This through-space mixing depends on the dipolar coupling which scales with the distance and orientation of the two spins being recoupled. From our MD simulations we note that due to steric considerations with the ring, each PIP headgroup phosphate has two main orientations, as monitored by the H-C-O-P dihedral angle. For both confirmations, the H-P distance to the nearest H is 2.8 Å. Thus, any changes in the HP CP efficiency of that transfer must be due to changes in the motions of the phosphate group resulting in averaging of the dipolar coupling or changes in T_1ρ_ relaxation times. Similarly, the interfacial glycerol phosphate can rotate relative to the nearest proton. Therefore, we decided to measure site specific HP CP transfer kinetics for each phosphate group in our samples by acquiring 2D HP spectra (Figure 4F) at a variety of ^1^H-^31^P CP mixing times. The intensity of each peak plotted against the contact time yields a characteristic “build up” curve that can be fit according to equation 1,^28^ which gives values for a CP time constant (T_IS_) a proxy for the dipolar coupling strength, and T_1ρ_, the rotating frame relaxation time.

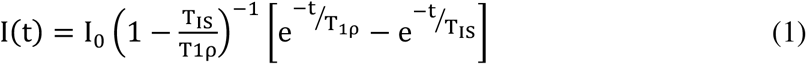

**Figure 4.**
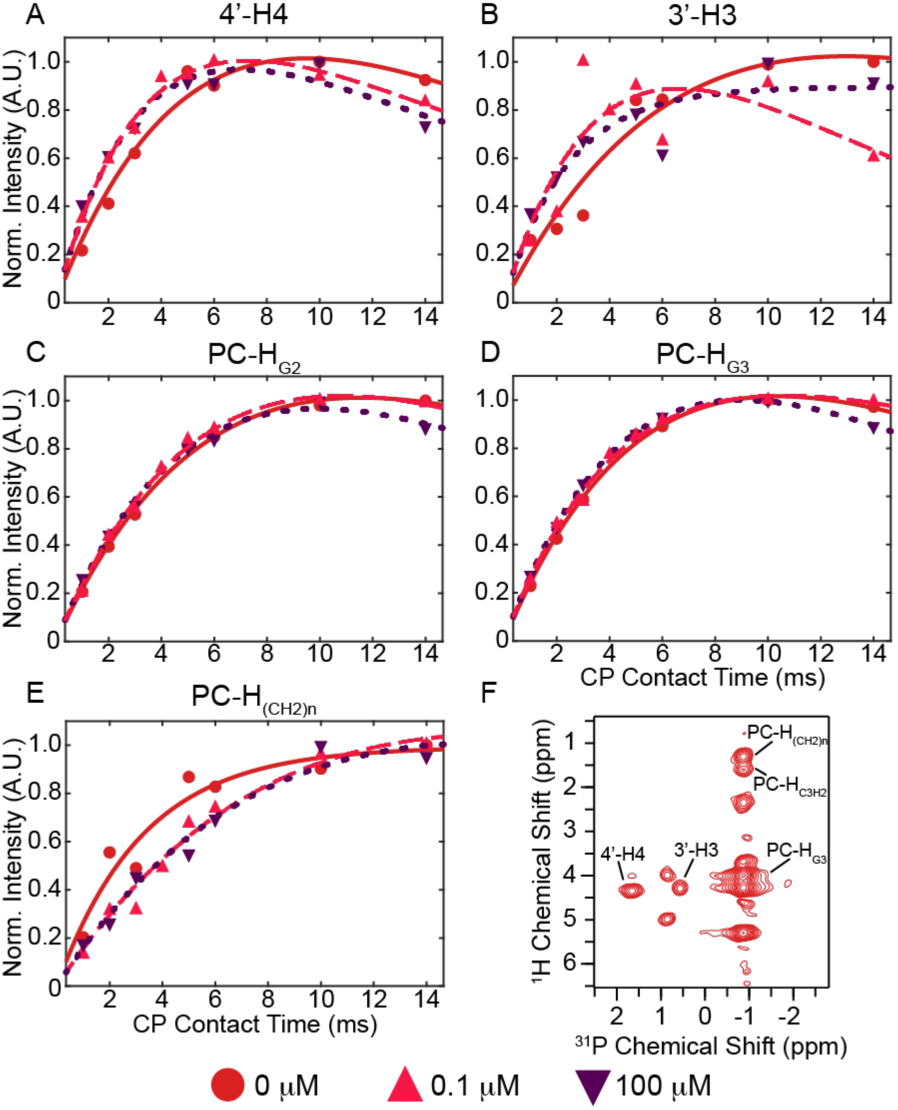
Cross polarization build ups for PIP3/PC at three different Ca^2+^ concentrations. Site specific CP build up curves are shown for ^1^H-^31^P correlations at important locations of the lipid. The A. 4’-H4 and B. 3’-H3 CP curves give insight into the head group of PIP3, while C. PC-H_G2_ and D. PC-H_G3_ are representative of the interfacial glycerol region, with emphasis on the effects on PC rather than PIP3, which is only 5% of the signal. CP curves were also generated for the tail region, specifically for E. PC-(CH_2_)_n_, which due to the distance between the PC glycerol backbone phosphate and the lipid tail protons, the signal of which is due to spin diffusion rather than cross polarization directly. F. The HP CP 2D with peak assignments is also shown.

HP CP build up curves were acquired at three different calcium concentrations for PIP3/PC liposomes across eight contact times ranging from 1-14 ms. The spectrum with no calcium (Figure 4F, S3A) shows the most ^31^P-^1^H correlation peaks in the PIP3 head group region as expected due to the longer T_2_ and sharper lines compared to the 0.1 μM (Figure S3B) and the 100 μM (Figure S3C) samples. The 0.1 μM showed the fewest cross peaks, further supporting our hypothesis of a more heterogeneous sample. Peak intensity for peaks that appear at all three Ca^2+^ concentrations decrease with increased Ca^2+^ concentration. We extracted the volume of each cross peak using the AutoFit routine in NMRPipe and fit the resulting data to equation 1 using MATLAB. The resulting curves are shown in Figure 4, plotted with the strongest peak in each normalized to 1 to allow for better comparisons of the shape of the curve. The fitting parameters can be found in the SI. The CP build up curves for the PIP3 head group ^31^P-^1^H correlation peaks for 4’-H4 (Figure 4A) and 3’- H3 (Figure 4B) both show similar trends with [Ca^2+^]. When analyzing these curves, we found that the 0 μM builds up the slowest with the 0.1 and 100 μM Ca^2+^ building up faster and beginning to build down by 14 ms CT. Given the lack of variation in the ^1^H-^31^P distance as discussed earlier, faster build up in the presence of calcium for the PIP3 head group peaks implies a decrease in motion, most likely due to structures like the ion mediated PIP3 dimer involving P3 and P4 shown in Figure 4E. Curves for PC-H_G2_ and PC-H_G3_ (Figure 4C and D) both show no significant change with Ca^2+^ present, meaning calcium is not affecting the overall rotation of the PC head group. These findings support the conclusion that PC is primarily not involved in the Ca^2+^-induced PIP3 clustering seen from the relaxation studies. The slower build up seen for the peak from the aliphatic tail CH_2_ to PC (Figure 4E) reflects the mixed nature of that transfer, with ^1^H-^1^H spin diffusion to nearby protons necessary prior to the ^1^H-^31^P CP transfer. Thus the slowing down of that transfer with Ca^2+^, despite no changes in the interfacial region (Figure 4C,D) the protons actually involved in CP, indicate that spin diffusion along the lipid tail slows in samples with Ca^2+^ present, evidence that heterogeneity at the headgroup level is also present deeper in the membrane.

With an understanding of how PIP3 behaves upon Ca^2+^ addition in a bilayer consisting of only zwitterionic PC, we now look to understand how the presence of another anionic lipid, PS, will alter this behavior. Looking again at the ^31^P 1D spectra, changes in chemical shifts of the inositol phosphates indicate a slight tendency for the anionic PS to deprotonate PIP3. The magnitude of this effect is slightly larger than was seen previously for PIP2 with 20% PS. PIP3 phosphate peaks have lower full width hall maximum (FWHM) for the 3’, 4’, and 5’ PIP peaks due to longer T_2_ which will be described in detail below. Figure 5A shows the ^31^P 1D spectra of PIP3/PC/PS liposomes at a series of Ca^2+^ concentrations. As with the PIP3/PC liposomes, the addition of even a small amount of Ca^2+^ results in decreased resolution and increased linewidth. In this case at 0.1 μM there are even broader lines which we attribute to this very small amount of calcium (1 per 100 PIP3 pairs) causing disorder due to populations with and without calcium. For 1 and 5 μM, the PIP3 head group phosphate peaks are more resolved than the PIP3/PC liposomes at the same [Ca^2+^], showing a combination of the broadening effect of the Ca and the narrowing effect observed with PS. The spectrum at 100 μM shows a significant decrease in resolution and peak intensity which we attribute to a similar situation as the 0.1 μM sample, with only 2 calcium for every 100 PS causing heterogeneity throughout the sample due to the tendency of PS to dimerize when bound to Ca^2+^. At 2 mM [Ca^2+^] this broadness has resolved, akin to the 100 μM for PIP3/PC without PS because each PS has at least one Ca^2+^ ion which decreases heterogeneity across the sample.

**Figure 5.**
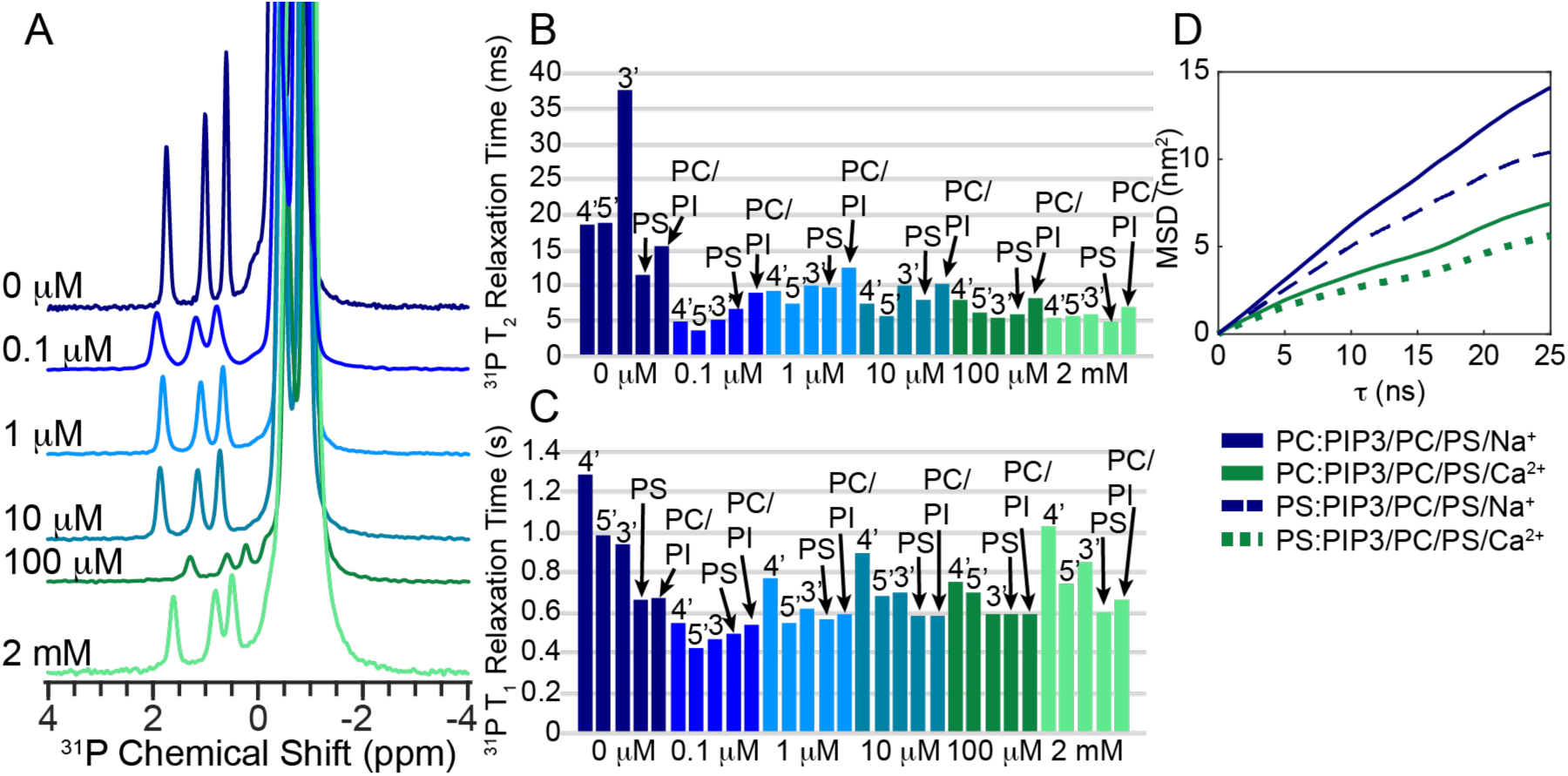
^31^P and MD simulations/lateral diffusion data for 5:70:25 PIP3/PC/PS liposomes at different Ca^2+^ concentrations from 0 μM-2 mM A. ^31^P 1D spectra of PIP3/PC/PS liposomes at each Ca^2+^ concentration. Changes in chemical shift, peak intensity, line width, and resolution are seen. B. Site specific ^31^P T_2_ relaxation data shows a large initial drop in T_2_ relaxation time for all phosphates on the PIP3 head group upon Ca^2+^ addition, which is also seen in the C. site specific ^31^P T_1_ relaxation data. D. The MD lateral diffusion data for PC within PIP3/PC/PS liposomes with Na^+^ (solid navy) and Ca^2+^ (solid green), and for PS within those same liposomes (dashed navy for Na^+^, dotted green for Ca^2+^).

For the dynamics experiments on the PIP3/PC/PS samples, the ^31^P T_2_ values (Figure 5B) show a significant decrease from 0 to 0.1 μM [Ca^2+^] but then increases for the 1 μM before continuing to decrease. We again attribute this initial rise to a decrease in heterogeneity following the very disordered first sample in the series. The ^31^P T_1_ values (Figure 5C) also show a significant decrease going from 0 to 0.1 μM then a gradual increase in ^31^P T_1_ values after that. Again, the decrease in ^31^P T_1_ and T_2_ observed by adding 0.1 μM Ca^2+^ indicates a decrease in the head group rotation of PIP3, PC, and PS in the presence of Ca^2+^. The falling T_2_/T_1_ ratio for these samples mirrors what was seen for the samples without PS. Again, this is indicative of an initial drop in rotational diffusion followed by a drop in lateral diffusion upon the addition of more calcium. The more gradual decrease of ^31^P T_2_ for the PIP3 head group phosphates differs from the PS-free PIP3/PC, as the PIP phosphates are in competition with the PS for Ca^2+^. Diffusion measurements from our HMMM MD simulations (Figure 5D) show that Ca^2+^ affects both PC and PS diffusion, with PC more affected in the presence of PS. While the addition of Ca^2+^ dropped PC diffusion by ∼30% in the PC/PIP3 membranes, both PS and PC show 45% decreases in diffusion upon the addition of Ca^2+^. This indicates that PC is not free to diffuse past larger PS clusters, perhaps due to PC-PS salt bridges. More evidence of the restriction of PC in the presence of PS will be seen in the CP build up curves in the following section.

HP CP 2D spectra were acquired at four different concentrations of PIP3/PC/PS across the same eight contact times ranging from 1-14 ms as the PIP3/PC liposomes (Figure S4). The HP 2D spectra of the PIP3 headgroup region show a slight decrease in ^31^P-^1^H correlation peak intensity at 0.1 μM Ca^2+^, but all of the same correlation peaks are still present. A significant decrease in both the number of correlation peaks present and the intensity of the remaining peaks is seen at 100 μM Ca^2+^. 2 mM Ca^2+^ exhibits even further loss of peaks and signal intensity than 100 μM. For the 4’- H4 PIP3 head group correlation peak (Figure 6A), no large change in CP build up is noted between the 0 and 0.1 μM [Ca^2+^], which indicates no significant Ca^2+^ effect on motion at the low concentration of 0.1 μM with PS present. The 100 μM and 2 mM [Ca^2+^] show faster build up for the 4’-H4, as well as faster build down. At these concentrations the PIP3 head group appears to have significant motional changes due to Ca^2+^ presence, as the 3’-H3 correlation peak (Figure 6B) shows significantly faster build up and subsequent faster build down for the 100 μM and 2 mM as well. The 3’-H3 peak does have a noticeable increase in build up for the 0.1 μM relative to the 0 μM, unlike the 4’-H4 peak, but it is still much less of an effect than is seen at the two higher Ca^2+^ concentrations. Comparing the build up behavior of these two PIP3 head group peaks in PIP3/PC/PS liposomes to the same peaks in PIP3/PC liposomes (Figure 5A and B), the decreased effect of 0.1 μM Ca^2+^ on the build up rates of these peaks with PS present is likely due to the low amount of Ca^2+^ being split between the PIP3 and PS head groups at this concentration, leaving many unbound and contributing to the sample heterogeneity seen in the ^31^P 1D spectrum (Figure 5A). For PC-H_G2_ (Figure 6C) with PS present, the PC-H_G2_ peak is now affected by the presence of calcium and shows a slightly slower build up rate and increase in motion (perhaps disorder) for 0.1 μM Ca^2+^ and higher build up rate and decrease in motion for 100 μM and 2 mM. Since we know that PC does not interact with Ca^2+^ directly, or changes would have been seen in the samples without PS, this change in headgroup rotational motion provides more evidence of PC-PS interactions. The PS correlation peak PS-H_G2_ shows the same slightly later peak in the build up curve for 0.1 μM Ca^2+^ and earlier peak for 100 μM and 2 mM. For this site, T_IS_ for all four [Ca^2+^] are not extremely different, rather T_1π_ is decreasing affecting the build down rate. The 0 μM builds down the slowest, the 2 mM the fastest, with the 0.1 μM and the 100 μM in between indicating decreased PS mobility at higher Ca^2+^.

**Figure 6.**
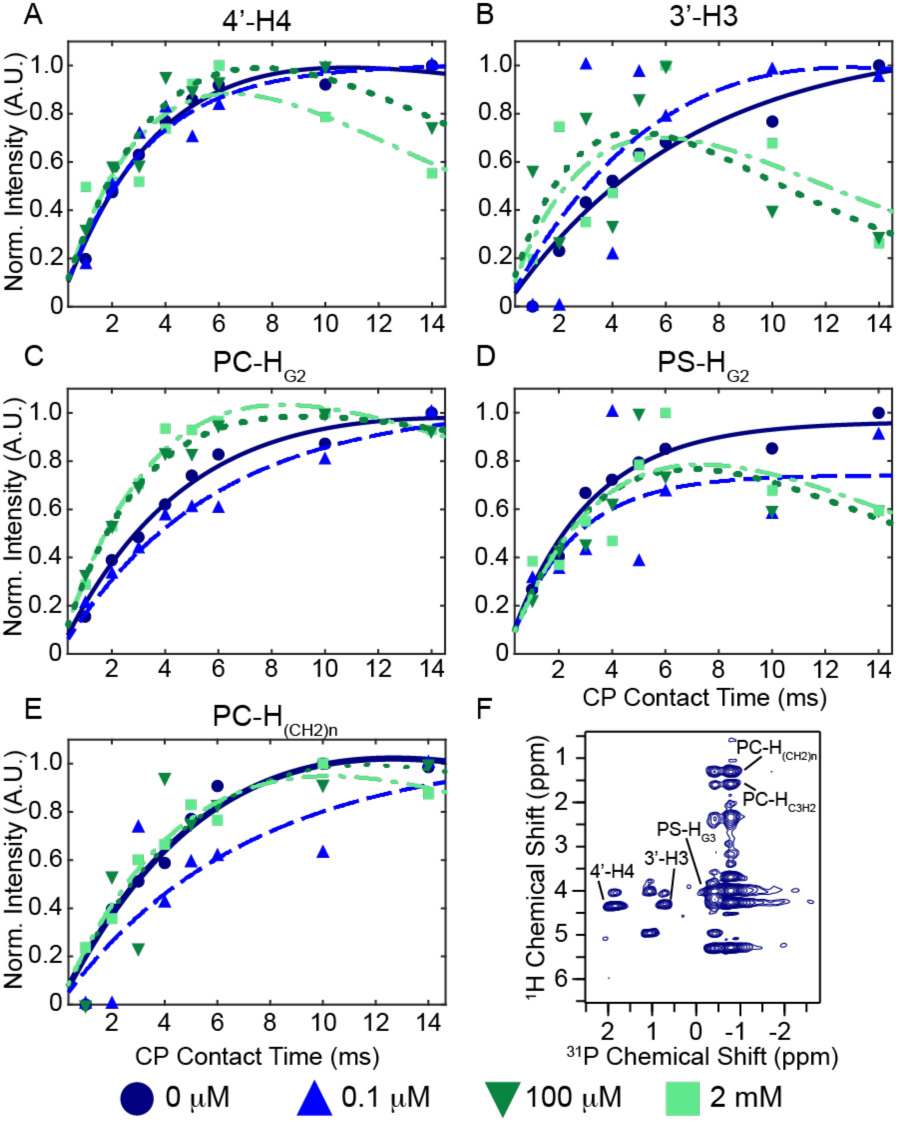
Cross polarization data of 5:70:25 PIP3/PC/PS liposomes at four different Ca^2+^ concentrations. Site specific CP build up curves are shown for H-P correlations at different regions of the lipid. The A. 4’-H4 and B. 3’-H3 CP curves give insight into the head group of PIP3, while C. PC-H_G2_ is representative of the interfacial glycerol region for PC. The interfacial region of PS D. PS-H_G2_ is also shown. CP curves were also generated for the tail region, specifically for E. PC- (CH_2_)_n_, which due to the distance between the PC glycerol backbone phosphate and the lipid tail protons, the signal of which is due to spin diffusion rather than cross polarization directly. F. The HP CP 2D with peak assignments is also shown.

**Figure 7.**
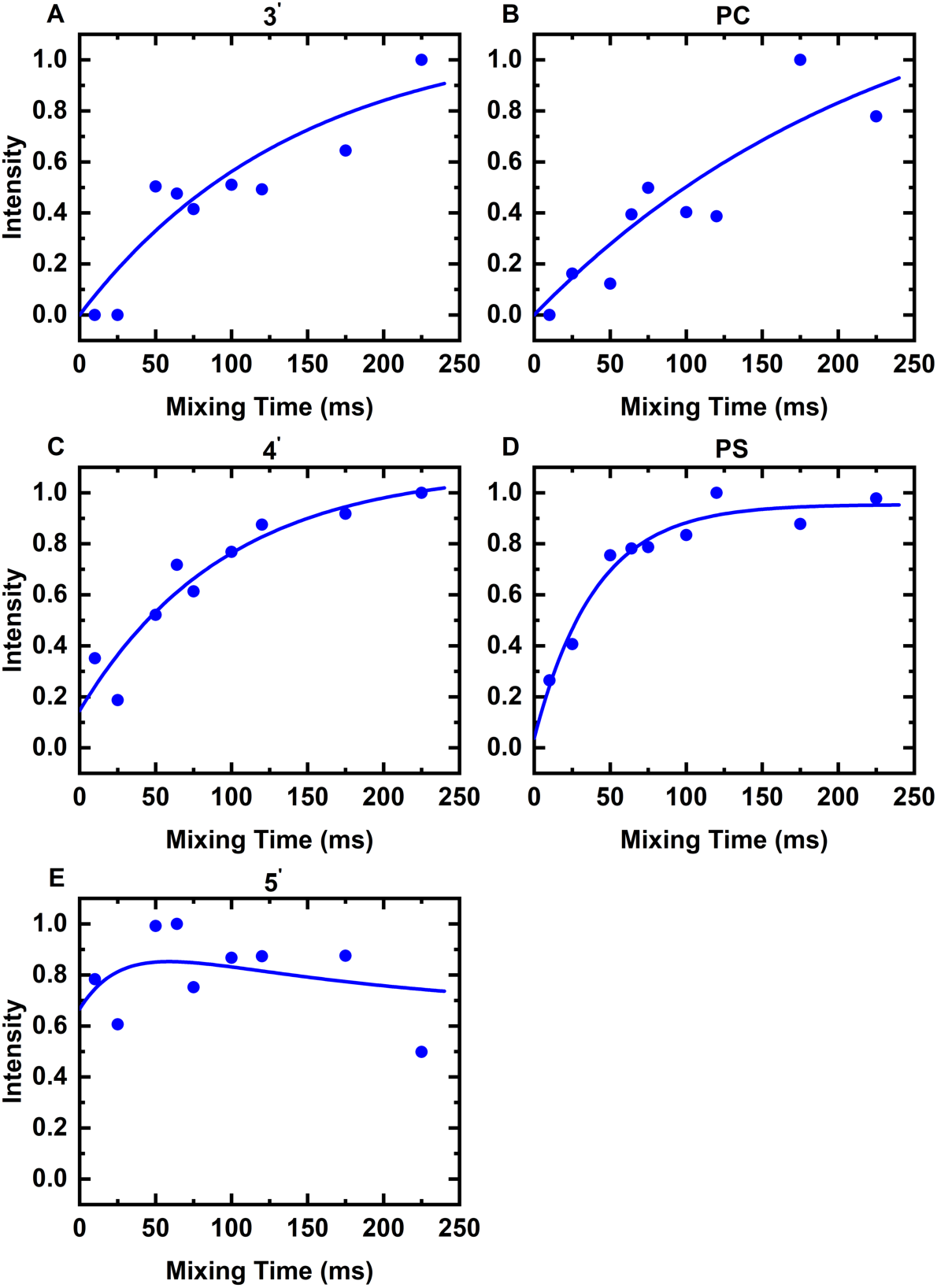
Proton-proton mixing kinetics for select H-P peaks after a water selective T_2_ filter. Magnetization built up for sites on the PIP headgroup (A, C, E) as well as in the interfacial region for PC (B) and PS (D) are shown. Volumes were extracted from 2D HP planes and fit to a biexponential equation (blue) to account for the possibility of mixing on to and off of a given site.

Up to this point we have refrained from discussing the HP 2D peaks associated with the 5’ phosphate group. Unlike the other two sites, the strongest peak for the 5’ does not correlate to the nearest proton on the inositol ring. Rather it occurs at 4.9 ppm, a chemical shift similar to that of tightly associated water. To test this possibility, we utilized a ^1^H T_2_ filter to eliminate signals from the lipids and only retain those from the solvent followed by a ^1^H-^1^H mixing period to allow signals to propagate back to the lipids, an experiment introduced by the lab of Mei Hong. The resulting spectra show only peaks to the 5’ and PS at the lowest mixing times, suggesting the presence of a bound water in exchange with the bulk at those two sites. Alternatively for the 5’ phosphate, one of the ^1^H on the phosphate itself may be in exchange with water. This would be consistent with the pKa determined without PS by Grienke^22^, although the pKas from same paper would indicate that similar exchange could be occurring for the 3’ and 4’. However, the need to be in exchange with bulk water in order to pick up the chemical shift might be the reason the 5’ site is most prominent, as the 3’ and 4’ sites are engaged in binding to ions preferentially in our MD simulations, while the 5’ is mostly engaged in a cross ring H-bond which might enhance the accessibility of the 5’ site to exchange with the bulk. Further evidence for a difference between the 5’ PIP and PS sites can be seen in the build up curves for the magnetization in the T_2_ filtered experiment seen in figure 8. The 5’ site (Figure 8E) shows maximal signal at the shortest mixing time, which might indicate the ^1^H is on the phosphate itself, while the PS (Figure 8D) shows a more gradual build up. Mixing to the 4’ (Figure 8C) is faster than the 3’ (Figure 8A), also consistent for with the exchangeable proton being on the 5’. We believe this indicates the transfer of signal through spin diffusion to the 3’ and 4’ phosphates and that they do not have exchangeable protons present bonded to them.

## CONCLUSIONS

Calcium titrations were carried out for PIP3-containing liposomes in the absence and presence of anionic lipid PS to characterize the effect of Ca^2+^ on PIP3 in lipid bilayers alone and with another anionic lipid. Calcium limits the head group mobility of PIP3 both with and without PS, though more calcium is needed to see the same effect as a lower concentration of Ca^2+^ without PS, as PS is also interacting with calcium, limiting the available Ca^2+^ for PIP3 to interact with. These interactions between calcium with both PIP3 and PS lead to clustering. We also see evidence of direct interactions between PC and PS. The clusters of anionic lipids and calcium decease lateral diffusion within the lipid bilayer, which is more pronounced with significant percentages of anionic lipid present in the lipid bilayer. This works demonstrates that significant amounts of information that can be obtained from unlabeled lipid systems by ssNMR.

## MATERIALS AND METHODS

### Liposome Preparation

All lipids were purchased from Avanti Polar Lipids, including POPC (1-palmitoyl-2-oleoyl-glycero-3-phosphocholine), POPS (1-palmitoyl-2-oleoyl-sn-glycero-3-phospho-L-serine), and PIP3 (1-stearoyl-2-arachidonoyl-sn-glycero-3-phospho-(1’-myo-inositol-3’,4’,5’-trisphosphate)). Chemicals used for buffer were obtained from Fisher Bioreagents (MES monohydrate and calcium chloride) and Sigma Aldrich (Sodium Chloride).

Lipid bilayers in the form of liposomes were prepared by the thin-film hydration method followed by sonication for 30 minutes in a bath sonicator.^29^ The buffer at pH 6.5 consisted of 50 mM MES, 50 mM NaCl, and a range of concentrations of CaCl_2_ as needed. The sonicated solutions were then pelleted in a Beckman Coulter Optima Max-TL ultracentrifuge at 100,000 rpm for 2-3 hours and packed into Phoenix 1.6 mm ssNMR rotors with a previously reported packing tool.^30^

### Solid-state NMR

All spectra were acquired on a Bruker Avance III HD spectrometer equipped with an HFXY 1.6 mm Phoenix probe tuned to HPC mode with frequencies of 599.54 MHz for ^1^H, 242.698 MHz for ^31^P, and 150.759 MHz for ^13^C. Spectra were acquired at 15 kHz and 20 °C (direct polarization spectra) and -2.5 °C (cross polarized spectra) unless otherwise mentioned. For the 20 °C spectra, the VTU was set to low, 600 lph flow, and 200 lph standby flow. The -2.5 °C chiller settings were strong, 600 lph flow, and 200 lph standby. 90° pulse lengths were as follows: 2.12 μs + 0.05 μs for ^1^H, 2.05 μs + 0.1 for ^31^P, and 2.04 μs + 0.02 μs for ^13^C. ^1^H 1D spectra were acquired with 4 scans and a recycle delay of 2.5 s. ^31^P 1D spinning spectra were acquired with 20000 or 4096 scans and a recycle delay of 2.5 s. Static ^31^P 1D spectra were acquired for at least 64 and up to 16834 scans with recycle delays of 3 or 4 s. HP CP 2D spectra were acquired for 128 or 64 or 32 scans in the direct dimension and either 256 or 196 rows in the indirect. HC 2D spectra were acquired in four blocks of 256 scans and 256 rows and added together in processing. Static ^31^P spectra can be found in figure S6.

### Solution NMR

Solution NMR experiments were performed on a 600.13 MHz Bruker Avance III HD spectrometer with an HCN probe. Dried lipids were suspended in deuterated buffer (50 mM each of NaCl and NaHPO_4_) at pH 6.5 and transferred into 5 mm NMR tubes. Spectra for PC/PS liposomes and IP4 were acquired at 40°C and 20°C respectively. Their respective ^1^H 90° pulse lengths were 9.44μs and 9.86μs. The ^1^H chemical shifts were referenced with DSS. HSQC 2D spectra were acquired for 512 scans in the direct dimension and 256 rows in the indirect dimension, with a recycle delay of 1.5s.

### Data Processing

All ^1^H spectra were referenced by setting the terminal methyl peak in the ^1^H 1D at 20 °C to 0.9 ppm, per literature.^31^ All ^31^P spectra were referenced to H_3_PO_4_,where the ^31^P chemical shift of H_3_PO_4_ was set to 0 ppm. ^13^C was referenced to adamantane. All 1D spectra and relaxation data were processed using TopSpin version 3.6.1 and MATLAB version R2020a. Excel also used in some cases. 2D spectra were processed in NMRPipe then analyzed in CCPN analysis version 2.4. NMRPipe was also used to add together 2D blocks. 2D CP build up curves were processed in NMRPipe then integrated using the Autofit protocol. Microsoft excel was then used to organize and normalize the data before utilizing MATLAB version R2020a and its curve fitting tool to fit the data to equation 10 from Kolodziejski et al.^28^ ChemDraw Professional 17.0 was used for drawing the chemical structures of the lipids. All figures were made using Adobe Illustrator 2020. <u>Dynamic Light Scattering</u>

Dynamic Light Scattering (DLS) was performed to determine average Rayleigh scattering radius for all liposome samples. A Wyatt Technology DynaPro Plate Reader equipped with DYNAMICS 7 software was used. Data was acquired at 25 °C with 5 scans per well, with each well consisting of 5 μL of liposomes pre-ultracentrifugation in 90 μL of whichever buffer they had been formed in. Microsoft excel was used for further analysis after data acquisition. All DLS data can be found in the supporting information (Figure S7 and Table S1).

### Molecular Dynamics Simulations

#### Preparation of membrane systems

All-atom molecular dynamics simulations were performed by employing (HMMM), which consist of short-tail lipids stuffed by 1,1-dichloroethane (DCLE) mimicking the hydrophobic core of the membrane cores. It has been suggested that the lateral diffusion of the short-tail lipids is 1-2 orders of magnitude faster than that of full-tail lipids. 95:5 PC:PIP3 and 70:25:5 PC:PS:PIP3 HMMM membranes were built using CHARMM-GUI in a box of initial size 8.5*8.5*10 nm^3^ wherein the membranes were hydrated with ∼13500 water molecules. One-microsecond simulation was performed for each membrane ionized with 400 mM Na^+^ and 200 mM Ca^2+^ respectively producing four 1-microsecond trajectories.

#### MD simulation protocols

MD simulations were performed using GROMACS-2021 package. Lipids and ions were modeled with CHARMM36 force field, and DCLE molecules with a CGenFFnforce field. Standard ion parameters without NBFIX pair-specific non-bond modifications were used to match better the ion solvation free energies. Water molecules were modeled explicitly using TIP3P parameters. Simulations were equilibrated with six simulations provided by CHARMM-GUI. The first two simulations were performed in the NVT ensemble with the magnitude of atomic restraints reduced after each simulation. The last four simulations were performed similarly in the NPT ensemble.

Equations of motion were integrated using the leapfrog algorithm with a 2-fs time step. The Nose-Hoover thermostat with a damping coefficient of 1 ps and the Parrinello-Rahman barostat with a damping coefficient of 5 ps were employed to hold a constant temperature of 310 K and a constant pressure of 1 atm for the simulations in NPT ensemble. A Verlet-list with van der Waals cutoff of 1.2 nm was employed to account for first neighbors. The smooth particle mesh Ewald scheme with a grid spacing of 0.12 nm and a real space cutoff of 1.2 nm were used to treat electrostatic interaction. A restraint of 20 kJ mol^-1^ nm^-2^ along the membrane normal was applied to the carboxy carbon atoms of each lipid to minimize the chance that the short-tail lipids travel into the solution. This simulation strategy does not impact the lateral motion of lipids and were demonstrated to be a good enhanced sampling approach in the study of membrane.

#### Analysis of MD simulations

The MD trajectories were analyzed using the built-in modules (i.e., *gmx msd, rotacf, rdf, and pairdist*) of GROMACS to extract lateral diffusion, rotational autocorrelation function, radial distribution, and distances. Dihedrals were computed using in-house Python codes with MDtraj package. Visualizations of molecules were rendered using VMD.

## AUTHOR INFORMATION

### Author Contributions

The manuscript was written through contributions of all authors. All authors have given approval to the final version of the manuscript.

### Funding Sources

This work was supported by the US National Institutes of Health (R01-GM139905 to A.J.N.).

## ACKNOWLEDGMENT

We thank Benjamin Lefkin for assistance with figures. We also would like to thank the NMR facility managers Seho Kim and Nagarajan Murali for their assistance. We acknowledge the Adam Gormley lab for the use of their DLS.

## ABBREVIATIONS

NMR: Nuclear Magnetic Resonance
ssNMR: Solid-state NMR
PIP3: Phosphatidylinositol Triphosphate
PS: Phosphatidylserine
PC: Phosphatidylcholine
MD: Molecular Dynamics

## Notes

### Competing Interest Statement

The authors have declared no competing interest.

## REFERENCES

1. Balla, T., Phosphoinositides: tiny lipids with giant impact on cell regulation. Physiol Rev 2013, 93 (3), 1019–137.

2. Schink, K. O.; Tan, K. W.; Stenmark, H., Phosphoinositides in Control of Membrane Dynamics. Annu. Rev. Cell Dev. Biol. 2016, 32, 143–171.

3. Di Paolo, G.; De Camilli, P., Phosphoinositides in cell regulation and membrane dynamics. Nature 2006, 443 (7112), 651–657.

4. Lemmon, M. A., Membrane recognition by phospholipid-binding domains. Nat. Rev. Mol. Cell Biol. 2008, 9 (2), 99–111.

5. van Meer, G.; de Kroon, A. I., Lipid map of the mammalian cell. J. Cell Sci. 2011, 124 (Pt 1), 5–8.

6. van Meer, G.; Voelker, D. R.; Feigenson, G. W., Membrane lipids: where they are and how they behave. Nat. Rev. Mol. Cell Biol. 2008, 9 (2), 112–24.

7. Ingolfsson, H. I.; Melo, M. N.; van Eerden, F. J.; Arnarez, C.; Lopez, C. A.; Wassenaar, T. A.; Periole, X.; de Vries, A. H.; Tieleman, D. P.; Marrink, S. J., Lipid Organization of the Plasma Membrane. J. Am. Chem. Soc. 2014, 136 (41), 14554–14559.

8. Manna, P.; Jain, S. K., Phosphatidylinositol-3,4,5-Triphosphate and Cellular Signaling: Implications for Obesity and Diabetes. Cell. Physiol. Biochem. 2015, 35 (4), 1253–1275.

9. Bagur, R.; Hajnoczky, G., Intracellular Ca2+ Sensing: Its Role in Calcium Homeostasis and Signaling. Mol. Cell 2017, 66 (6), 780–788.

10. Therrien, C.; Di Fulvio, S.; Pickles, S.; Sinnreich, M., Characterization of Lipid Binding Specificities of Dysferlin C2 Domains Reveals Novel Interactions with Phosphoinositides. Biochemistry 2009, 48 (11), 2377–2384.

11. Evans, J. H.; Gerber, S. H.; Murray, D.; Leslie, C. C., The calcium binding loops of the cytosolic phospholipase A(2) C2 domain specify targeting to Golgi and ER in live cells. Mol. Biol. Cell 2004, 15 (1), 371–383.

12. Ellenbroek, W. G.; Wang, Y. H.; Christian, D. A.; Discher, D. E.; Janmey, P. A.; Liu, A. J., Divalent Cation-Dependent Formation of Electrostatic PIP2 Clusters in Lipid Monolayers. Biophys. J. 2011, 101 (9), 2178–2184.

13. Brown, D. A., PIP2Clustering: From model membranes to cells. Chem. Phys. Lipids 2015, 192, 33–40.

14. Graber, Z. T.; Wang, W.; Singh, G.; Kuzmenko, I.; Vaknin, D.; Kooijman, E. E., Competitive cation binding to phosphatidylinositol-4,5-bisphosphate domains revealed by X-ray fluorescence. Rsc Adv 2015, 5 (129), 106536–106542.

15. Levental, I.; Christian, D. A.; Wang, Y. H.; Madara, J. J.; Discher, D. E.; Janmey, P. A., Calcium-Dependent Lateral Organization in Phosphatidylinositol 4,5-Bisphosphate (PIP2)- and Cholesterol-Containing Monolayers. Biochemistry 2009, 48 (34), 8241–8248.

16. Palmere, R. D.; Case, D. A.; Nieuwkoop, A. J., Simulations of Kindlin-2 PIP binding domains reveal protonation-dependent membrane binding modes. Biophys. J. 2021, 120 (24), 5504–5512.

17. Huang, B. X.; Akbar, M.; Kevala, K.; Kim, H. Y., Phosphatidylserine is a critical modulator for Akt activation. J. Cell Biol. 2011, 192 (6), 979–992.

18. Tuttle, M. D.; Comellas, G.; Nieuwkoop, A. J.; Covell, D. J.; Berthold, D. A.; Kloepper, K. D.; Courtney, J. M.; Kim, J. K.; Barclay, A. M.; Kendall, A.; Wan, W.; Stubbs, G.; Schwieters, C. D.; Lee, V. M.; George, J. M.; Rienstra, C. M., Solid-state NMR structure of a pathogenic fibril of full-length human alpha-synuclein. Nat. Struct. Mol. Biol. 2016, 23 (5), 409–15.

19. Retel, J. S.; Nieuwkoop, A. J.; Hiller, M.; Higman, V. A.; Barbet-Massin, E.; Stanek, J.; Andreas, L. B.; Franks, W. T.; van Rossum, B. J.; Vinothkumar, K. R.; Handel, L.; de Palma, G. G.; Bardiaux, B.; Pintacuda, G.; Emsley, L.; Kuhlbrandt, W.; Oschkinat, H., Structure of outer membrane protein G in lipid bilayers. Nat Commun 2017, 8 (1), 2073.

20. Boettcher, J. M.; Davis-Harrison, R. L.; Clay, M. C.; Nieuwkoop, A. J.; Ohkubo, Y. Z.; Tajkhorshid, E.; Morrissey, J. H.; Rienstra, C. M., Atomic view of calcium-induced clustering of phosphatidylserine in mixed lipid bilayers. Biochemistry 2011, 50 (12), 2264–73.

21. Doherty, T.; Hong, M., 2D H-1-P-31 solid-state NMR studies of the dependence of inter-bilayer water dynamics on lipid headgroup structure and membrane peptides. J. Magn. Reson. 2009, 196 (1), 39–47.

22. Kooijman, E. E.; King, K. E.; Gangoda, M.; Gericke, A., Ionization properties of phosphatidylinositol polyphosphates in mixed model membranes. Biochemistry 2009, 48 (40), 9360–71.

23. Akbarzadeh, A.; Rezaei-Sadabady, R.; Davaran, S.; Joo, S. W.; Zarghami, N.; Hanifehpour, Y.; Samiei, M.; Kouhi, M.; Nejati-Koshki, K., Liposome: classification, preparation, and applications. Nanoscale Res Lett 2013, 8.

24. Sturm, L.; Ulrih, N. P., Basic Methods for Preparation of Liposomes and Studying Their Interactions with Different Compounds, with the Emphasis on Polyphenols. International Journal of Molecular Sciences 2021, 22 (12).

25. Graber, Z. T.; Jiang, Z. P.; Gericke, A.; Kooijman, E. E., Phosphatidylinositol-4,5-bisphosphate ionization and domain formation in the presence of lipids with hydrogen bond donor capabilities. Chem. Phys. Lipids 2012, 165 (6), 696–704.

26. Lindon, J. C.; Baker, D. J.; Williams, J. M.; Irvine, R. F., Confirmation of the Identities of Inositol 1,3,4-Trisphosphate and Inositol 1,3,4,5-Tetrakisphosphate by the Use of One-Dimensional and Two-Dimensional Nmr-Spectroscopy. Biochem. J. 1987, 244 (3), 591–595.

27. Graber, Z. T.; Gericke, A.; Kooijman, E. E., Phosphatidylinositol-4,5-bisphosphate ionization in the presence of cholesterol, calcium or magnesium ions. Chem. Phys. Lipids 2014, 182, 62–72.

28. Kolodziejski, W.; Klinowski, J., Kinetics of cross-polarization in solid-state NMR: A guide for chemists. Chem Rev 2002, 102 (3), 613–628.

29. Bangham, A. D.; Standish, M. M.; Watkins, J. C., Diffusion of univalent ions across the lamellae of swollen phospholipids. J. Mol. Biol. 1965, 13 (1), 238–52.

30. Hisao, G. S.; Harland, M. A.; Brown, R. A.; Berthold, D. A.; Wilson, T. E.; Rienstra, C. M., An efficient method and device for transfer of semisolid materials into solid-state NMR spectroscopy rotors. J Magn Reson 2016, 265, 172–6.

31. Volke, F.; Pampel, A., Membrane Hydration and Structure on a Subnanometer Scale as Seen by High-Resolution Solid-State Nuclear-Magnetic-Resonance - Popc and Popc/C(12)Eo(4) Model Membranes. Biophys. J. 1995, 68 (5), 1960–1965.

