## Supplemental Information for "Effects of Ca^2+^ on the Structure and Dynamics of PIP3 in Model Membranes Containing PC and PS"

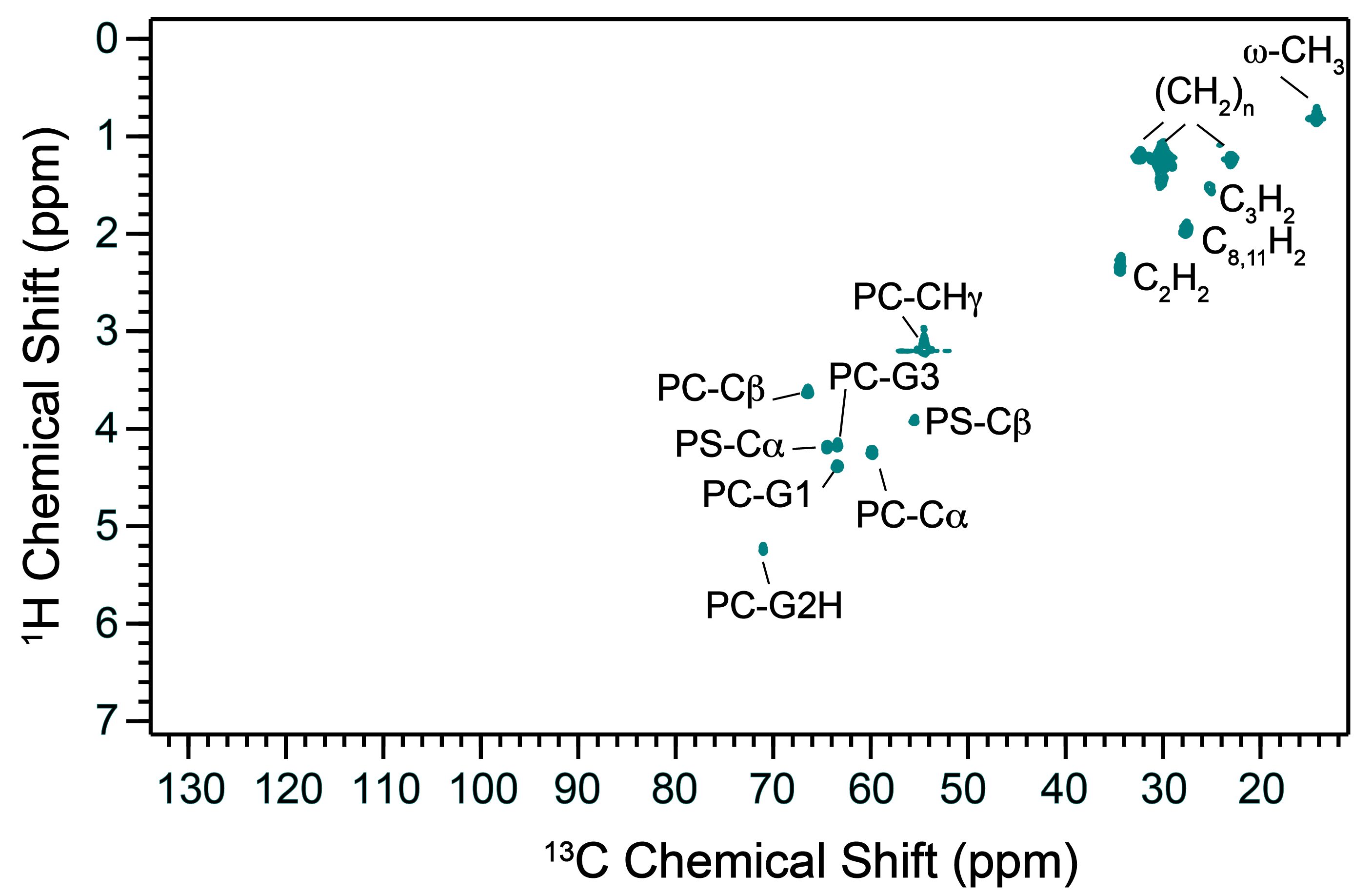

**Figure S1.** Solution HSQC spectrum of 70:30 PC/PS liposomes acquired at 40 °C and 600 MHz.

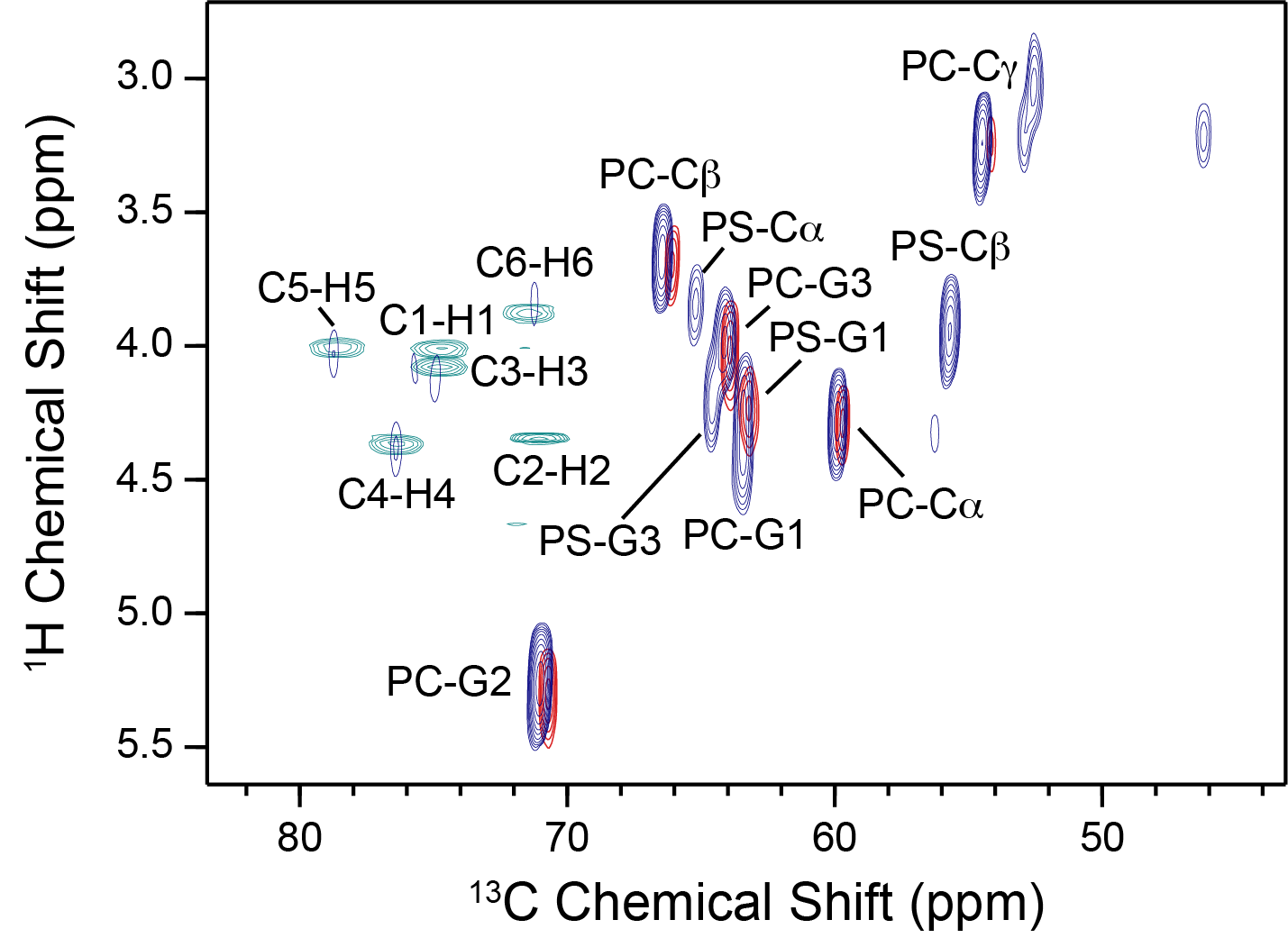

**Figure S2.** Overlay of solution HSQC of soluble PIP3 head group analogue Inositol 1,3,4,5-tetrakisphosphate (IP4) (teal) with solid-state HETCOR of PIP3/PC liposomes (red) and solid-state INEPT of PIP3/PC/PS liposomes (navy).

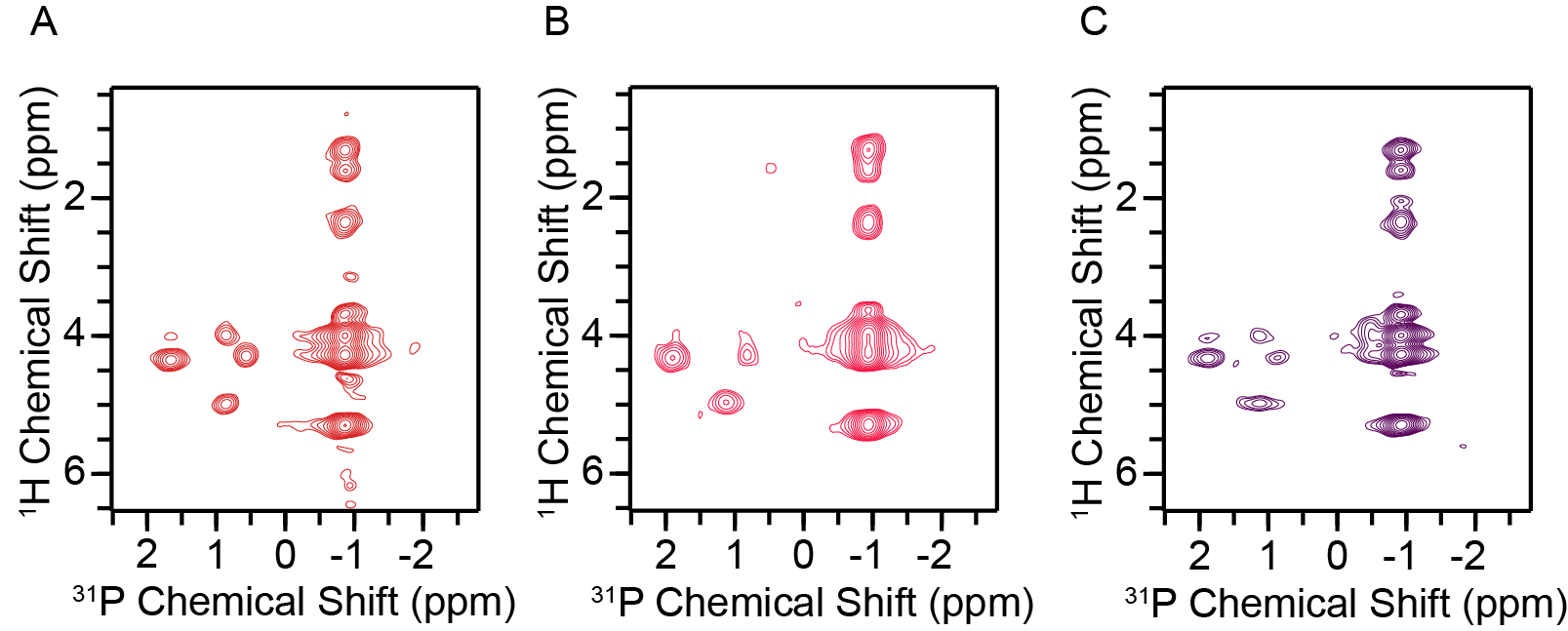

**Figure S3.** HP 2D of PIP3/PC with A. 0 μM Ca^2+^, B. 0.1 μM Ca^2+^, and C. 100 μM Ca^2+^. All spectra acquired at -2.5 °C and 15 kHz at 600 MHz.

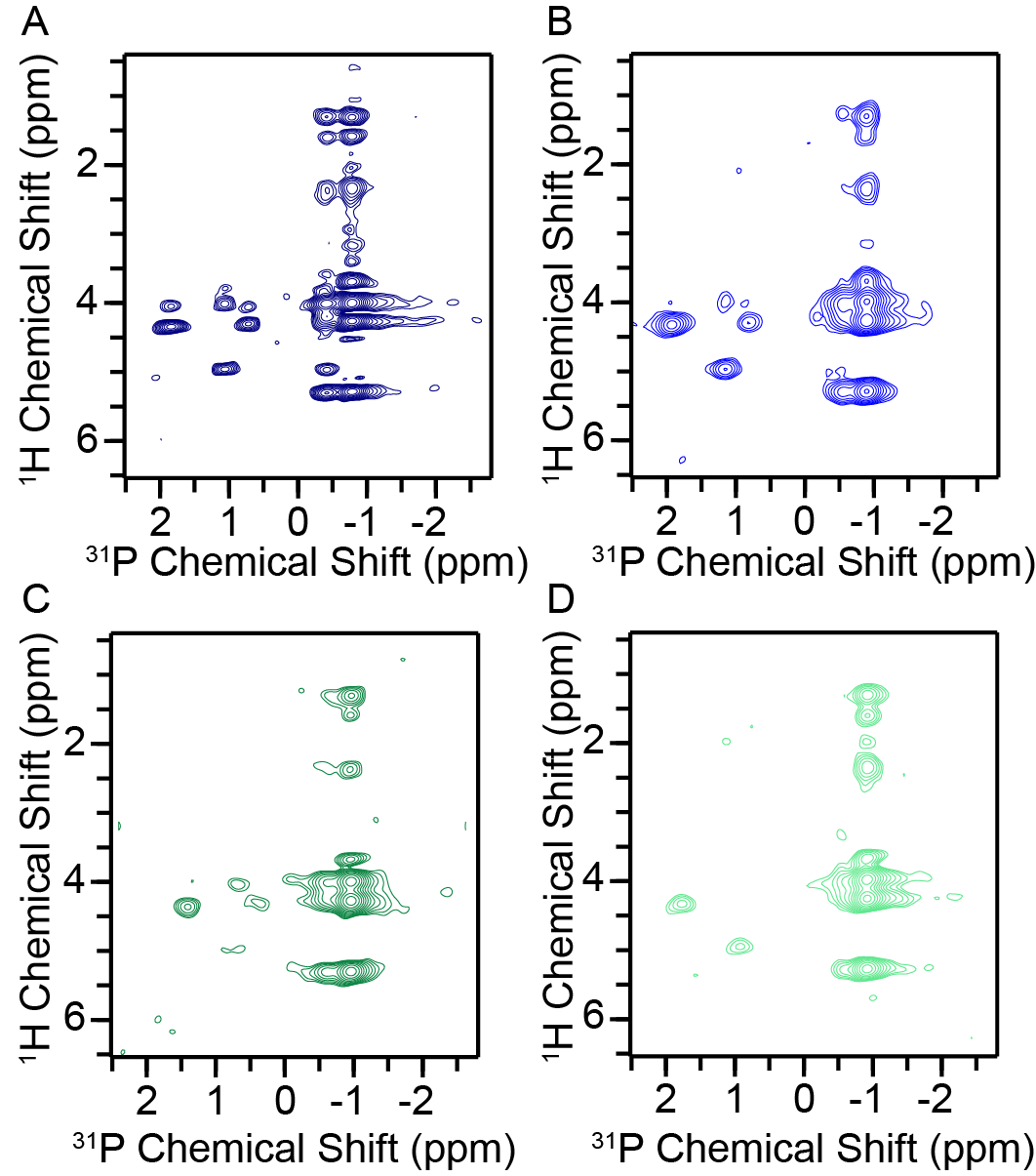

**Figure S4.** HP 2D of PIP3/PC/PS with A. 0 μM Ca^2+^, B. 0.1 μM Ca^2+^, C. 100 μM Ca^2+^, and D. 2 mM Ca^2+^. All spectra acquired at -2.5 °C and 15 kHz at 600 MHz.

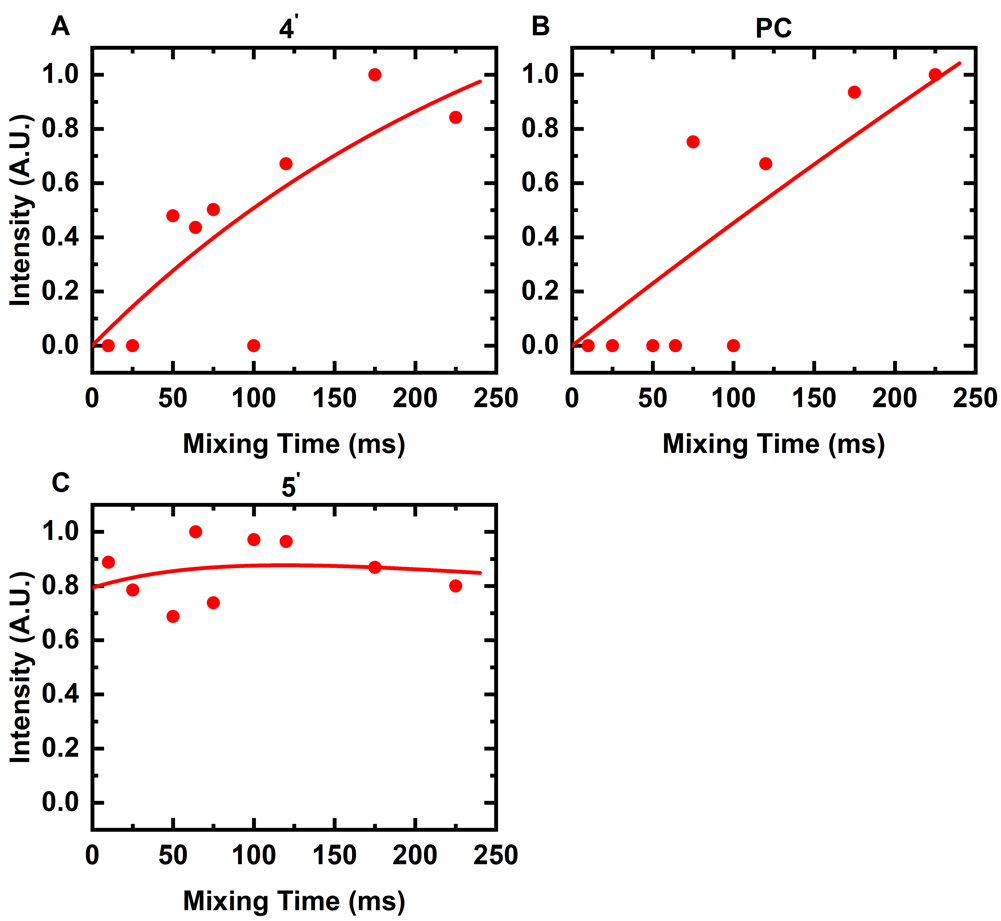

**Figure S5.** Proton-proton mixing kinetics for select ^1^H-^31^P peaks after a water selective T2 filter for PIP3/PC liposomes. Magnetization built up for sites on the PIP headgroup (A, C) as well as in the interfacial region for PC (B) are shown. Volumes were extracted from 2D HP planes and fit to a biexponential equation (red) to account for the possibility of mixing on to and off of a given site.

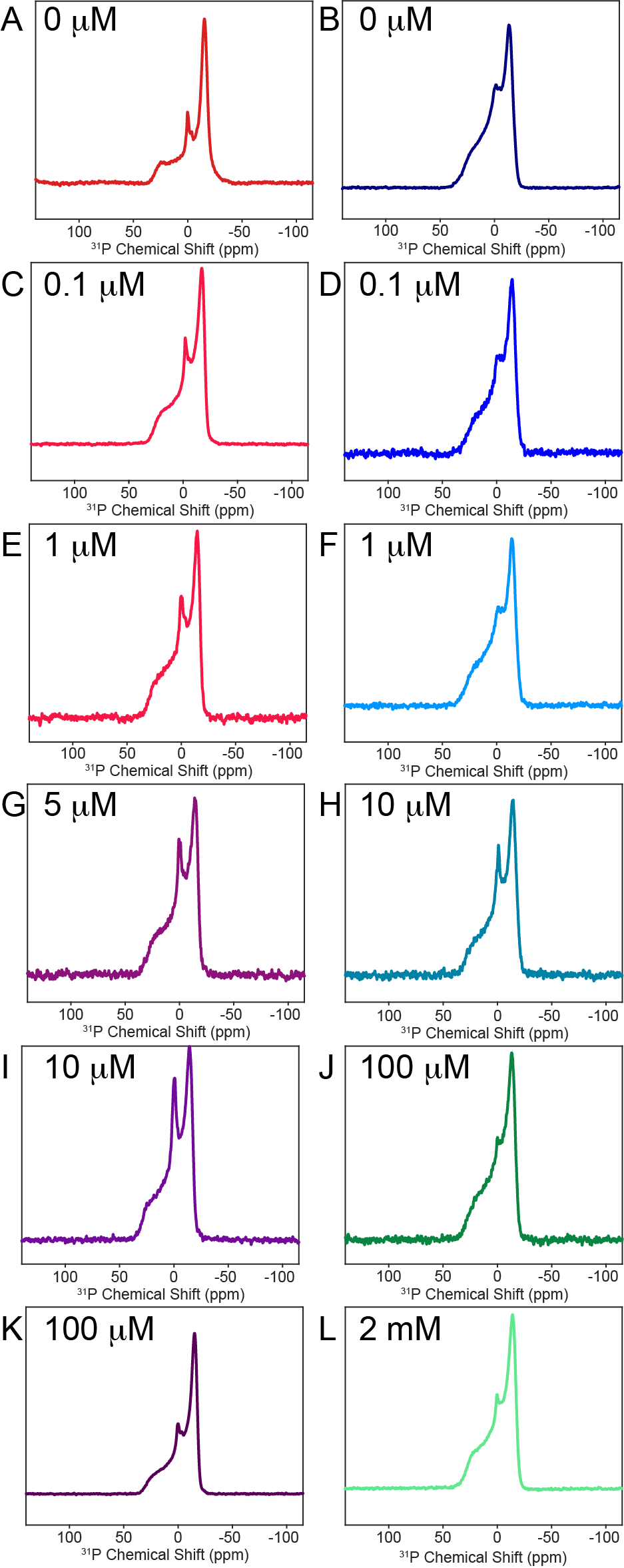

**Figure S6.** ^31^P Static 1D spectra for the Ca^2+^ titration for PIP3/PC liposomes (A, C, E, G, I, K) and PIP3/PC/PS liposomes (B, D, F, H, J, L) at each Ca^2+^ concentration.

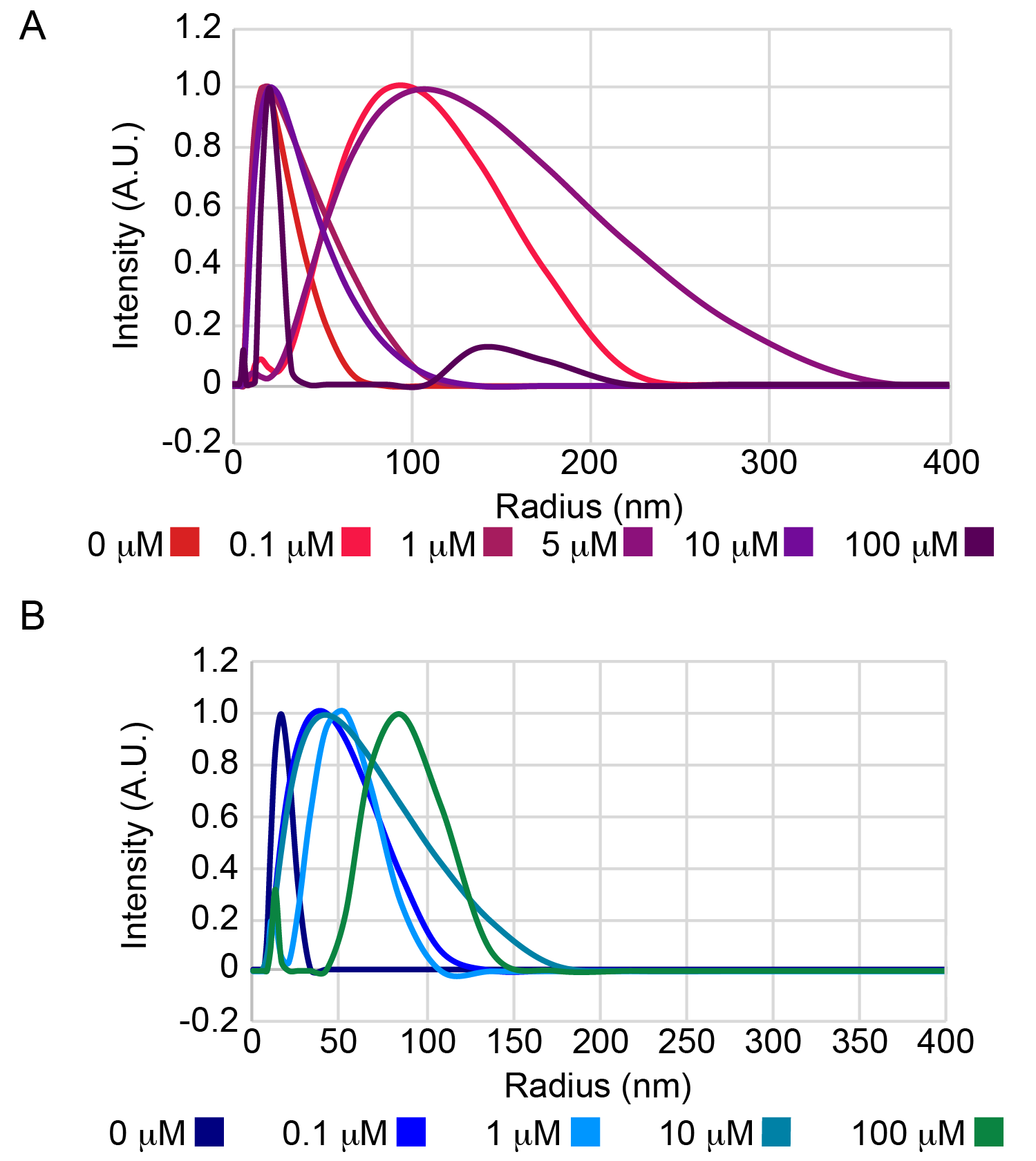

**Figure S7.** DLS size distributions for Ca^2+^ titration samples of A. PIP3/PC and B. PIP3/PC/PS.

**Table S1.** DLS determined Radii for PIP3-containing liposomes at different [Ca^2+^]

| **Sample Composition** | **[Ca^2+^] μM** | **Hydrodynamic radius (nm)** | **Sample Composition** | **[Ca^2+^] μM** | **Hydrodynamic radius (nm)** |
| --- | --- | --- | --- | --- | --- |
| PIP3/PC | 0 | 18.7 | PIP3/PC/PS | 0 | 17.2 |
|  | 0.1 | 76.9 |  | 0.1 | 32.6 |
|  | 1 | 20.4 |  | 1 | 37.7 |
|  | 5 | 89.6 |  | 10 | 32.8 |
|  | 10 | 21.5 |  | 100 | 66.9 |
|  | 100 | 20.7 |  |  |  |
